# A chromosome-scale high-contiguity genome assembly of the threatened cheetah (*Acinonyx jubatus*)

**DOI:** 10.1101/2022.12.16.520814

**Authors:** Sven Winter, René Meißner, Carola Greve, Alexander Ben Hamadou, Petr Horin, Stefan Prost, Pamela A. Burger

## Abstract

The cheetah (*Acinonyx jubatus*, SCHREBER 1775) is a large felid and is considered the fastest land animal. Historically, it inhabited open grassland across Africa, the Arabian Peninsula, and southwestern Asia; however, only small and fragmented populations remain today. Here, we present a de novo genome assembly of the cheetah based on PacBio continuous long reads and Hi-C proximity ligation data. The final assembly (*VMU_Ajub_asm*_v1.0) has a total length of 2.38 Gb, of which 99.7% are anchored into the expected 19 chromosome-scale scaffolds. The contig and scaffold N50 values of 96.8 Mb and 144.4 Mb, respectively, a BUSCO completeness of 95.4% and a k-mer completeness of 98.4%, emphasize the high quality of the assembly. Furthermore, annotation of the assembly identified 23,622 genes and a repeat content of 40.4%. This new highly contiguous and chromosome-scale assembly will greatly benefit conservation and evolutionary genomic analyses and will be a valuable resource, e.g., to gain a detailed understanding of the function and diversity of immune response genes in felids.

## Introduction

The cheetah (*Acinonyx jubatus*, SCHREBER 1775) is a large carnivore of the cat family Felidae, in which it forms the tribe Acinonychini together with the puma (*Puma concolor*), and the jaguarundi (*Herpailurus yagouaroundi*) (Durant et al., 2021). The cheetah is known as the fastest land animal, as it reaches speeds of up to 105 km/h (Sharp, 1997). Historically, it occurred in open grasslands across Africa, the Arabian Peninsula, and southwestern Asia (Durant et al., 2017). At present, it only inhabits small fractions of its former range resulting in small and fragmented populations (Durant et al., 2017). The cheetah, as a species, is currently considered “vulnerable” on the International Union for Conservation of Nature (IUCN) Red List of threatened species, with two subspecies *A. j. venaticus* (Iran) and *A. j. hecki* (Northwest Africa), being listed as “critically endangered” (Belbachir, 2008; Durant et al., 2021; Farhadinia et al., 2017). The use and need for genomic analyses to support conservation decisions and management has increasingly been recognized, e. g., by the IUCN Cat Specialist Group (IUCN Cat Specialist Group, 2021). Recently, conservation and evolutionary genomic analyses of the cheetah based on short-read sequences have been published (Dobrynin et al., 2015; Prost et al., 2022). However, in-depth conservation or evolutionary genomic analyses, such as the inference of runs of homozygosity or analyses of mutational load and genetic health, greatly benefit from highly continuous reference genomes currently unavailable for the cheetah. Thus, we sequenced and assembled a chromosome-level genome for the cheetah with a much-improved continuity than previously reported genome assemblies.

## Material and Methods

### Sequencing and Assembly

High molecular weight genomic DNA was extracted from the blood of a 14-year-old female cheetah named “Pintada” (GAN:27869175) from Lisbon Zoo, Portugal, using the PacBio Nanobind CBB kit (PacBio, Menlo Park, CA, USA). The blood was drawn during routine veterinary procedures and immediately frozen at -20°C. DNA concentration and molecule length were evaluated using the Qubit dsDNA BR Assay kit on the Qubit Fluorometer (Thermo Fisher Scientific) and the Genomic DNA Screen Tape on the Agilent 2200 TapeStation system (Agilent Technologies), respectively.

Two sequencing libraries were prepared, one PacBio continuous long read (CLR) library using the SMRTbell Express Prep Kit 2.0 (PacBio) and one short read library using the NEBNext Ultra II FS DNA Library Kit for Illumina (New England BioLabs Inc., Ipswich, MA, USA). The long-read library was then sequenced on the PacBio Sequel IIe system in CLR mode using the Sequel II Binding Kit 2.2 (PacBio). The short-read library with an insert size of 350 bp was sequenced on the NovaSeq6000 platform (Illumina, Inc., San Diego, CA, USA), generating 150 bp paired-end reads.

After receiving the data from the Sequel IIe run, we converted the PacBio subreads from BAM to FASTQ format using BAM2fastx v.1.3.0, a PacBio Secondary Analysis Tool (https://github.com/PacificBiosciences/pbbioconda). We then used Flye v.2.9 (<u>RRID:SCR_017016</u>) (Kolmogorov et al., 2019) to assemble the reads into a contig-level genome assembly. Flye was run with the options for raw PacBio reads and the default parameters, including one iteration of long-read polishing.

The short-read data were first trimmed using fastp v.0.20.1 (<u>RRID:SCR_016962</u>) (Chen et al., 2018) with base correction and low complexity filter enabled to remove sequencing adaptors and polyG stretches at the end of reads. We also employed a 4 bp sliding window to detect regions of poor quality (Phred score <15). Reads were removed if they fit into one of the following categories: reads below 36 bp length, reads with >40% low-quality bases, or reads with 5 or more undetermined bases (Ns). The trimmed reads were then mapped to the assembly using bwa-mem v. 0.7.17 (<u>RRID:SCR_010910</u>) (Li, 2013). The resulting mapping file was then sorted by position in the assembly, converted to BAM format, and indexed using samtools v.1.9 (<u>RRID:SCR_002105</u>) (Li et al., 2009). The mapped short reads were used to further improve the base-level accuracy of the assembly with one iteration of short-read polishing with pilon v.1.23 (<u>RRID: SCR_014731</u>) (Walker et al., 2014).

To anchor the polished contigs into chromosome-scale scaffolds, we utilized previously generated proximity ligation (Hi-C) data for the same subspecies from the DNAZoo (www.dnazoo.org, accession numbers: SRR8616936, SRR8616937). First, we followed the Arima Hi-C mapping pipeline used by the Vertebrate Genome Project (https://github.com/VGP/vgp-assembly/blob/master/pipeline/salsa/arima_mapping_pipeline.sh) for filtering and mapping of the data to the assembly. In short, the pipeline mapped the reads to the assembly using bwa-mem v.0.7.17 (<u>RRID:SCR_010910</u>) (Li, 2013) and filtered the mapped reads with samtools v.1.14 (<u>RRID:SCR_002105</u>) (Li et al., 2009) based on multiple parameters such as mapping quality, read quality, CIGAR strings, etc. Subsequently, duplicated reads were marked and removed using the Picard v. 2.26.10 (<u>RRID:SCR_006525</u>) (Broad Institute, 2019) tool “MarkDuplicates”. The mapped and filtered reads were then used in YaHS v.1.1 (<u>RRID:SCR_022965</u>) (Zhou et al., 2022) for proximity-ligation-based scaffolding. Hi-C contact maps were generated with JuicerTools v. 1.22.01 (<u>RRID:SCR_017226</u>) (Durand et al., 2016) and used for manual curation of the scaffolded assembly in Juicebox v.1.11.08 (<u>RRID:SCR_021172</u>) (Durand et al., 2016). Furthermore, we run TGS-GapCloser v.1.1.1 (<u>RRID:SCR_017633</u>) (Xu et al., 2020) for two iterations to close gaps in the assembly and increase its contiguity. Each iteration of gap-closing utilized a random subset of approximately 25% of the long-read data to reduce computational requirements, as well as the short reads for polishing. The long-read subsets were generated from the complete dataset with seqtk v. 1.3 (<u>RRID:SCR_018927</u>) (Li, 2018a) using the random number generator seeds 11 and 18.

To evaluate the quality and completeness of the assembly, we generated assembly statistics with Quast v.5.0.2 (<u>RRID:SCR_001228</u>) (Gurevich et al., 2013), ran a gene set completeness analysis with BUSCO v.5.3.1 (<u>RRID:SCR_015008</u>) (Manni et al., 2021) using the carnivora_odb10 dataset and compared the results to the previously available chromosome-scale assembly from DNAZoo (Dobrynin et al., 2015; Dudchenko et al., 2017), which is based on an earlier draft genome (Dobrynin et al., 2015). We also evaluated the mapping rate of both the short and long reads with QualiMap v.2.2.1 (<u>RRID:SCR_001209</u>) (Okonechnikov et al., 2016) after mapping the reads to our assembly with bwa-men v.0.7.17 (<u>RRID:SCR_010910</u>) (Li, 2013) and minimap2 v.2.17 (<u>RRID:SCR_018550</u>) (Li, 2018b), respectively. In addition, we analyzed the completeness, base-level error rate, and quality value (QV) of the assembly based on a k-mer size of 21 using Merqury v.1.1 (<u>RRID:SCR_022964</u>) (Rhie et al., 2020). We further evaluated potential contamination with BlobTools v.3.2.4 (<u>RRID:SCR_017618</u>) (Laetsch & Blaxter, 2017) utilizing the generated mapping files and a BLASTN v.2.11.0+ (<u>RRID:SCR_001598</u>) (Zhang et al., 2000) search against the NCBI Nucleotide database.

In addition to the nuclear genome, we also assembled the mitochondrial genome from the short reads with GetOrganelle v1.7.5 (<u>RRID:SCR_022963</u>) (Jin et al., 2020).

### Annotation

For increased accuracy during gene prediction, repeat regions in the assembly were first masked. RepeatModeler v. 2.0.1 (<u>RRID:SCR_015027</u>) (Flynn et al., 2020) was used to generate a de novo repeat library, which was then combined with the Felidae repeat dataset

(July 2022) from RepBase (<u>RRID:SCR_021169</u>) (Bao et al., 2015). This custom repeat library was then used to annotate and mask the repeats in the genome using RepeatMasker v.4.1.0 (http://www.repeatmasker.org/RMDownload.html, RRID:SCR_012954). We hard-masked all interspersed repeats and soft-masked simple repeats.

Genes were predicted using the homology-based gene prediction with MMseqs2 (<u>RRID:SCR_022962</u>) (Steinegger & Söding, 2017) as alignment tool implemented in the GeMoMa pipeline v.1.7.1 (<u>RRID:SCR_017646</u>) (Keilwagen et al., 2016, 2018). We used the following nine mammalian genomes and associated annotations as references: Homo sapiens (GCF_000001405.40), *Mus musculus* (GCF_000001635.27), *Lynx canadensis* (GCF_007474595.2), *Canis lupus familiaris* (GCF_014441545.1), *Prionailuris bengalensis* (GCF_016509475.1), *Leopardus geoffroyi* (GCF_018350155.1), *Felis catus* (GCF_018350175.1), *Panthera tigris* (GCF_018350195.1), and *Panthera leo* (GCF_018350215.1).

Functional annotation of the predicted proteins was conducted by a BLASTP v.2.11.0+ (<u>RRID:SCR_001010</u>) (Zhang et al., 2000) search with an e-value cutoff of 10^−6^ against the Swiss-Prot database (<u>RRID:SCR_002380</u>; release 2021-02). Furthermore, we annotated gene ontology (GO) terms, domains, and motifs using InterProScan v.5.50.84 (<u>RRID:SCR_005829</u>) (Jones et al., 2014; Quevillon et al., 2005). The completeness of the predicted proteins was evaluated with BUSCO v.5.3.1 (<u>RRID:SCR_015008</u>) (Manni et al., 2021) using the *carnivora_odb10* dataset.

### Synteny between feline genomes

We analyzed synteny between *VMU_Ajub_asm_v1*.*0*, the previously published DNAZoo cheetah assembly (Dobrynin et al., 2015; Dudchenko et al., 2017), the tiger *Panthera tigris* (GCF_018350195.1), two assemblies of the domestic cat *Felis catus* (Fca126: GCF_018350175.1, Bredemeyer et al., 2021; Fca9.1: GCA_000181335.5), the leopard cat *Prionailurus bengalensis* (GCA_016509475.2; Bredemeyer et al., 2021), and the Canadian lynx *Lynx canadensis* (GCA_007474595.2; Rhie et al., 2021) using JupiterPlot v.3.8.1 (<u>RRID:SCR_022961</u>) (Chu, 2018). The closely related leopard cat was used as a reference to identify homologous cheetah chromosomes, as the chromosome structure based on G-banding is very similar to the cheetah (O’Brien et al., 2006).

## Results and Discussion

### Genome sequencing and assembly

Sequencing generated 341 Gb of long-read PacBio data or approximately 136-fold coverage with a mean subread length of 10,308.5 bp and approximately 18-fold (45.5 Gb) of short-read Illumina data.

After assembly with Flye, polishing with pilon, proximity-ligation scaffolding with YaHS, and two iterations of gap-closing, the final assembly (*VMU_Ajub_asm_v1*.*0*) had a total length of 2.38 Gb in 198 scaffolds (including the mitochondrial genome) and a scaffold and contig N50 of 144.4 Mb and 96.8 Mb, respectively (Table 1A). The largest 19 scaffolds (> 40 Mb), representing the expected haploid chromosome number of the cheetah (2n = 38) (O’Brien et al., 2006), span 99.7% of the total assembly length, resulting in a scaffold L50 of seven. *VMU_Ajub_asm_v1*.*0* is highly contiguous and reflects a major improvement in contiguity compared to the previously available cheetah genome assembly from DNA Zoo (Dobrynin et al., 2015; Dudchenko et al., 2017), as evidenced by a more than 3,000-fold larger contig NG50 (96.8 Mb vs. 32.2 kb) (Table 1A). The high quality and completeness of *VMU_Ajub_asm_v1*.*0* were also highlighted by a BUSCO completeness score of 95.4%, an increase of 5.7% compared to the previously available assembly, and Merqury k-mer-based completeness of 98.4%, with an error rate of 0.013% (QV = 38.7).

**Table 1.** Assembly statistics of *VMU_Ajub_asm_v1*.*0* in comparison to the previously available DNAZoo assembly (A) and repeat content of *VMU_Ajub_asm_v1*.*0* (B)

|  | Scaffold-level |  | Contig-level |  |
| --- | --- | --- | --- | --- |
|  | *VMU_Ajub_asm_v1.0* | DNAZoo | *VMU_Ajub_asm_v1.0* * | DNAZoo* |
| **No. of scaffolds/contigs** | 198 | 13,047 | 220 | 163,014 |
| **No. of scaffolds/contigs (> 1 kbp)** | 197 | 2,247 | 219 | 130,184 |
| **L50** | 7 | 7 | 9 | 19,040 |
| **LG50<sup>#</sup>** | 7 | 7 | 9 | 21,646 |
| **N50 (bp)** | 144,444,042 | 144,637,309 | 96,827,784 | 35,115 |
| **NG50<sup>#</sup> (bp)** | 144,444,042 | 144,637,309 | 96,827,784 | 32,192 |
| **Max. scaffold/contig length (bp)** | 235,669,126 | 235,519,777 | 218,150,482 | 419,587 |
| **Total length (bp)** | 2,377,450,114 | 2,373,338,770 | 2,377,445,714 | 2,328,406,444 |
| **GC (%)** | 41.59 | 41.29 | 41.59 | 41.29 |
| **No. of N's** | 4400 | 42,858,800 | 0 | 128,625 |
| **No. of N's per 100 kbp** | 0.19 | 1807.45 | 0 | 5.52 |

\*broken into contigs at gaps with a length of $\geq 10$ N's. Statistics for these columns are based on contigs, the remaining columns are based on scaffolds.
<sup>#</sup>based on an estimated reference length of 2,503,680,000 bp calculated from a C-value of 2.56 pg (genomesize.com)

**B**
| Type of element | Number of elements | Length (bp) | Percentage of assembly |
| --- | --- | --- | --- |
| **SINEs** | 887,856 | 129,629,255 | 5.45% |
| **LINEs:** | 1,755,353 | 586,984,696 | 24.69% |
| **L1/LINE1** | 1,558,262 | 535,696,713 | 22.53% |
| **LTR elements** | 371,149 | 128,445,131 | 5.40% |
| **DNA transposons** | 384,830 | 73,338,402 | 3.08% |
| Unclassified | 10,953 | 2,596,934 | 0.11% |
| Small RNA | 612,496 | 89,904,390 | 3.78% |
| Satellites | 3,697 | 921,773 | 0.04% |
| Simple repeats | 748,674 | 32,528,191 | 1.37% |
| Low complexity | 91,753 | 5,217,093 | 0.22% |
| Total | 4,868,672 | 960,526,883 | 40.40% |

### Annotation

#### Repeat annotation

A repeat content of 40.4% or 960.5 Mb of the sequence of *VMU_Ajub_asm_v1*.*0* was identified by the repeat annotation (Table 1B). Long Interspersed Nuclear Elements (LINEs) were the most abundant, spanning nearly one-quarter (24,7%) of the genome, followed by Short Interspersed Nuclear Elements (SINEs) and Long Terminal Repeat (LTR) Elements with 5.45% and 5.4%, respectively. The remaining repeat classes, such as DNA Transposons, small RNA, simple repeats, etc., each accounted for less than 4%.

#### Gene annotation

The homology-based gene prediction with GeMoMa identified 23,622 genes in *VMU_Ajub_asm_v1*.*0* with a median gene length of 7,857.5 bp spanning 468.4 Mb of the total assembly length. A BUSCO score of 89.2% of identified complete Carnivora orthologous genes suggest high annotation completeness (Figure 1B). InterProScan functionally annotated 66,775 out of the 67,405 predicted proteins (99.1%) and assigned at least one Gene Ontology (GO) term to 50,735 proteins (75,3%). In addition, more than 96.9% (65,333) of the predicted proteins were identified from the Swiss-Prot database.

**Figure 1.**
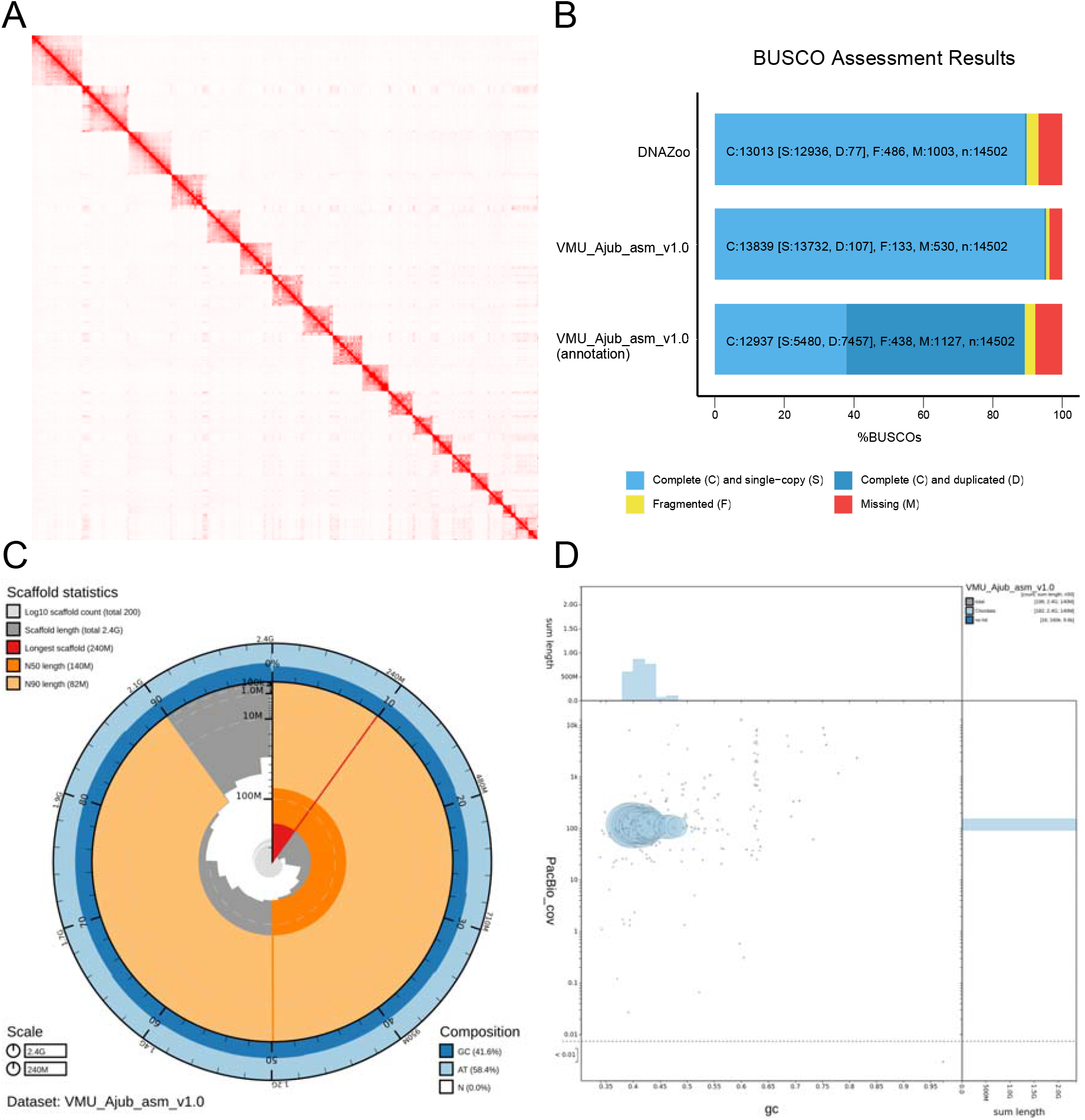
Assembly quality assessment of *VMU_Ajub_asm_v1*.*0*. (A) Hi-C contact density map depicting the 19 distinct chromosome-level scaffolds. (B) BUSCO gene set completeness analyses for the assembly, annotation (predicted proteins), and previously available DNAZoo assembly for comparison. (C) SnailPlot summarizing assembly statistics. (D) BlobPlot analysis comparing GC content (x-axis), sequencing depth of PacBio reads (y-axis), and taxonomic assignment of contigs (colors) show no evidence of contamination.

#### Synteny between feline genomes

JupiterPlots showed high levels of synteny between *VMU_Ajub_asm_v1*.*0* and other felid species, as expected by the identical chromosome numbers (2n=38) and the conserved nature of Felidae genomes (O’Brien et al., 2006) (Figure 2). Therefore, we were able to assign chromosome names to the 19 chromosome-scale scaffolds of *VMU_Ajub_asm_v1*.*0* according to the karyotype format commonly used for felids (O’Brien et al., 2006). We based the naming on the homologous regions of the leopard cat genome (Figure 2C), whose chromosomes have very similar G-banding patterns as the cheetah chromosomes (O’Brien et al., 2006; Wurster-Hill & Gray, 1973). Comparing *VMU_Ajub_asm_v1*.*0* and the previous DNAZoo assembly (Figure 2A) found structural differences only in the form of one translocation in chromosome D2 (scaffold 14) and an inversion in the smallest chromosome E4 (scaffold 19). However, both differences could potentially be scaffolding or assembly errors in one of the assemblies, despite both utilizing Hi-C data for scaffolding from the same individual. The most structural differences are evident between VMU_Ajub_asm_v1.0 and the most recent domestic cat assembly (Fca126, GCF_018350175.1, Figure 2D). Yet, when compared to the previous cat assembly (Fca9.1, GCA_000181335.5, Figure 2E), only very small rearrangements were identified, suggesting potential assembly errors in the latest cat assembly.

**Figure 2.**
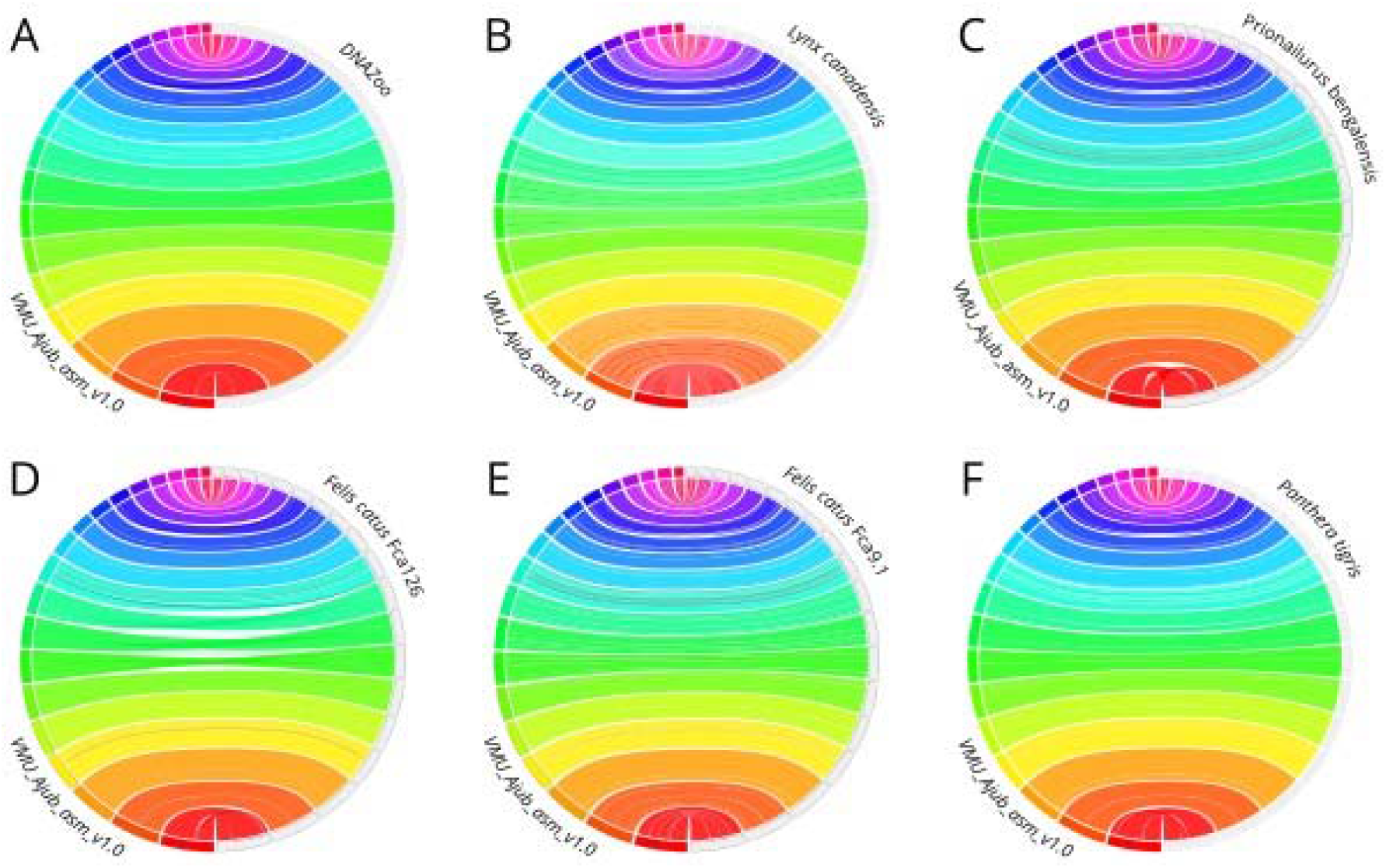
Synteny between the cheetah *Acinonyx jubatus* and other felid species. Circos plots generated with JupiterPlot comparing the synteny of the chromosome-scale cheetah genome assembly *VMU_Ajub_asm_v1*.*0* (A-F, left, colored boxes) with six available chromosome-scale assemblies of other felids (right, gray boxes): (A) A previous cheetah assembly from DNA Zoo, (B) the Canadian lynx *Lynx canadensis*, (C) the leopard cat *Prionailurus bengalensis*, (D&E) the domestic cat Felis catus (Fca126, Fca9.1), and (F) the tiger Panthera tigris. Colored ribbons between scaffolds indicate syntenic regions. Chromosome-scale scaffolds are sorted by size from the largest (bottom) to the smallest (top).

## Conclusion

Highly contiguous annotated chromosome-scale genome assemblies are valuable references for evolutionary or conservation genomic analyses and enable in-depth studies on structural variation, or the diversity and function of certain genes (e.g., immune response genes). However, genome assemblies from non-model organisms of this quality are still relatively rare. The presented new cheetah assembly *VMU_Ajub_asm_v1*.*0*, which is the first long-read-based assembly for this species, has a much-improved contiguity and will thus enable more in-depth genomic analyses for this threatened species. This genome resource provides a solid foundation to address key biological questions like understanding the process of natural selection and adaptation.

## Acknowledgments

We thank the Genome Technology Center (RGTC) at Radboudumc for the use of the Sequencing Core Facility (Nijmegen, The Netherlands), which provided the PacBio SMRT sequencing service on the Sequel IIe platform. We also thank Rui Bernardino from the zoo of Lisbon for the cheetah sample.

